# Subtyping evaluation of *Salmonella* Enteritidis using SNP and core genome MLST with nanopore reads

**DOI:** 10.1101/2022.05.03.490560

**Authors:** Zhihan Xian, Shaoting Li, David Ames Mann, Yixiao Huang, Feng Xu, Xingwen Wu, Silin Tang, Guangtao Zhang, Abigail Stevenson, Chongtao Ge, Xiangyu Deng

## Abstract

Whole genome sequencing (WGS) for public health surveillance and epidemiological investigation of foodborne pathogens predominantly relies on sequencing platforms that generate short reads. Continuous improvement of long-read nanopore sequencing such as Oxford Nanopore Technologies (ONT) presents a potential for leveraging multiple advantages of the technology in public health and food industry settings, including rapid turnaround and onsite applicability in addition to superior read length. However, evaluation, standardization and implementation of the ONT approach to WGS-based, strain-level subtyping is challenging, in part due to its relatively high base-calling error rates and frequent iterations of sequencing chemistry and bioinformatic analytics. Using an established cohort of *Salmonella* Enteritidis isolates for subtyping evaluation, we assessed the technical readiness of ONT for single nucleotide polymorphism (SNP) analysis and core-genome multilocus sequence typing (cgMLST) of a major foodborne pathogen. By multiplexing three isolates per flow cell, we generated sufficient sequencing depths under seven hours of sequencing for robust subtyping. SNP calls by ONT and Illumina reads were highly concordant despite homopolymer errors in ONT reads (R9.4.1 chemistry). *In silico* correction of such errors allowed accurate allelic calling for cgMLST and allelic difference measurements to facilitate heuristic detection of outbreak isolates. Our study established a baseline for the continuously evolving nanopore technology as a viable solution to high quality subtyping of *Salmonella*, delivering comparable subtyping performance when used standalone or together with short-read platforms.

## INTRODUCTION

Whole genome sequencing (WGS)-based subtyping and characterization of bacterial isolates has become a routine practice in foodborne disease surveillance and investigation, ushering in a paradigm shift in public health microbiology often referred to as genomic epidemiology (1–3). In public health laboratories, WGS of foodborne pathogens is predominantly performed on sequencing platforms that generate short reads up to a few hundred base pairs in length (4). After WGS, the short reads are typically used for genome-wide subtyping including single nucleotide polymorphisms (SNP) analysis and core genome multilocus sequence typing (cgMLST) (5, 6) as well as *in silico* characterizations such as antibiotic resistance (7) and serotype (8) profiling. While short reads are generally adequate for most of these tasks, they often do not yield gap-free and complete bacterial genomes, which facilitate SNP calling by serving as high-quality reference genomes and reduce missing cgMLST allelic calls caused by assembly gaps.

Recent and continuous improvement of long-read nanopore sequencing such as Oxford nanopore technologies (ONT) has created substantial interest in its evaluation for genomic epidemiology applications. In addition to facilitating genome assembly by producing long reads averaging 10-30 kilobases (kb), nanopore sequencing is available on portable devices and amenable for real-time data analysis, making it appealing for scenarios that require rapid turnaround (9) or onsite applicability (10).

Prior evaluation studies of ONT have generated promising yet mixed results. Nanopore sequencing has been shown reliable for determining common serotypes (11) and less common serotype variants (12) of *Salmonella enterica*, even using multiplexed sequencing that combined multiple isolates in a single sequencing run (13). Long ONT reads outperform short Illumina reads when the contiguity of genome assembly matters for genotyping analyses such as plasmid and virulence factor profiling (14). While short Illumina reads and long ONT reads were reported concordant enough for inferring SNP phylogeny of Shiga toxin-producing *Escherichia coli* (15), a similar comparative study using *Salmonella enterica* concluded that ONT reads alone were still insufficient for phylogenetic and pangenome analyses (16).

The discrepancy was in part rooted in elevated base calling errors associated with ONT reads (17), which affect different subtyping and profiling analyses to different degrees. A particular base calling challenge for ONT reads is the determination of the correct size of homopolymeric tracts, which has prompted empirical correction of homopolymer errors for bacterial multilocus sequence typing in the traditional seven housekeeping gene format (18). Another confounding factor is the constantly evolving sequencing chemistry and analytical algorithms of ONT reads. In contrast with the matured implementation of Illumina sequencing for foodborne pathogen WGS (19), the still highly dynamic development of nanopore sequencing complicates and challenges evaluation, standardization, and implementation of ONT platforms in public health surveillance and investigation of foodborne pathogens where stable and standardized methodology is often favored.

The most commonly adopted WGS subtyping methods for strain-level differentiation of bacterial pathogens are SNP and cgMLST (5). The latter approach has yet to be rigorously evaluated with ONT reads. The higher base calling errors in ONT reads are potentially problematic for cgMLST because correct cgMLST sequence type (ST) identification requires accurate sequence determination of thousands of loci across bacterial genomes. Previous evaluations of SNP subtyping of major foodborne pathogens using ONT reads were relatively limited in scale, including two clinical isolates from a potential outbreak (15) or non-outbreak human and food isolates of unclear epidemiological link (16).

In this study, we aimed to assess the technical readiness of using ONT reads for strain-level subtyping with an expanded scale and epidemiological representation. We analyzed a documented set of *Salmonella enterica* serotype Enteritids isolates, which represented 15 major outbreaks from 2001 to 2012 in the United States. This isolate set was previously used for comparative benchmarking of molecular subtyping against the WGS standard (20). We evaluated the R9.4.1 sequencing chemistry for SNP typing and cgMLST analysis against the benchmark of Illumina-ONT hybrid assemblies. We further demonstrated the possibility of robust cgMLST even with the evaluated sequencing chemistry that generated relatively frequent yet empirically correctable errors in homopolymeric tracts.

## MATERIALS AND METHODS

### Bacterial isolates

Fifty *S*. Enteritidis isolates previously analyzed for a comparative evaluation of molecular subtyping methods (20) were re-sequenced using Illumina and ONT platforms. These isolates were obtained from the National *Salmonella* Reference Laboratory at the Centers for Diseases Control and Prevention (Table 1).

**Table 1.**
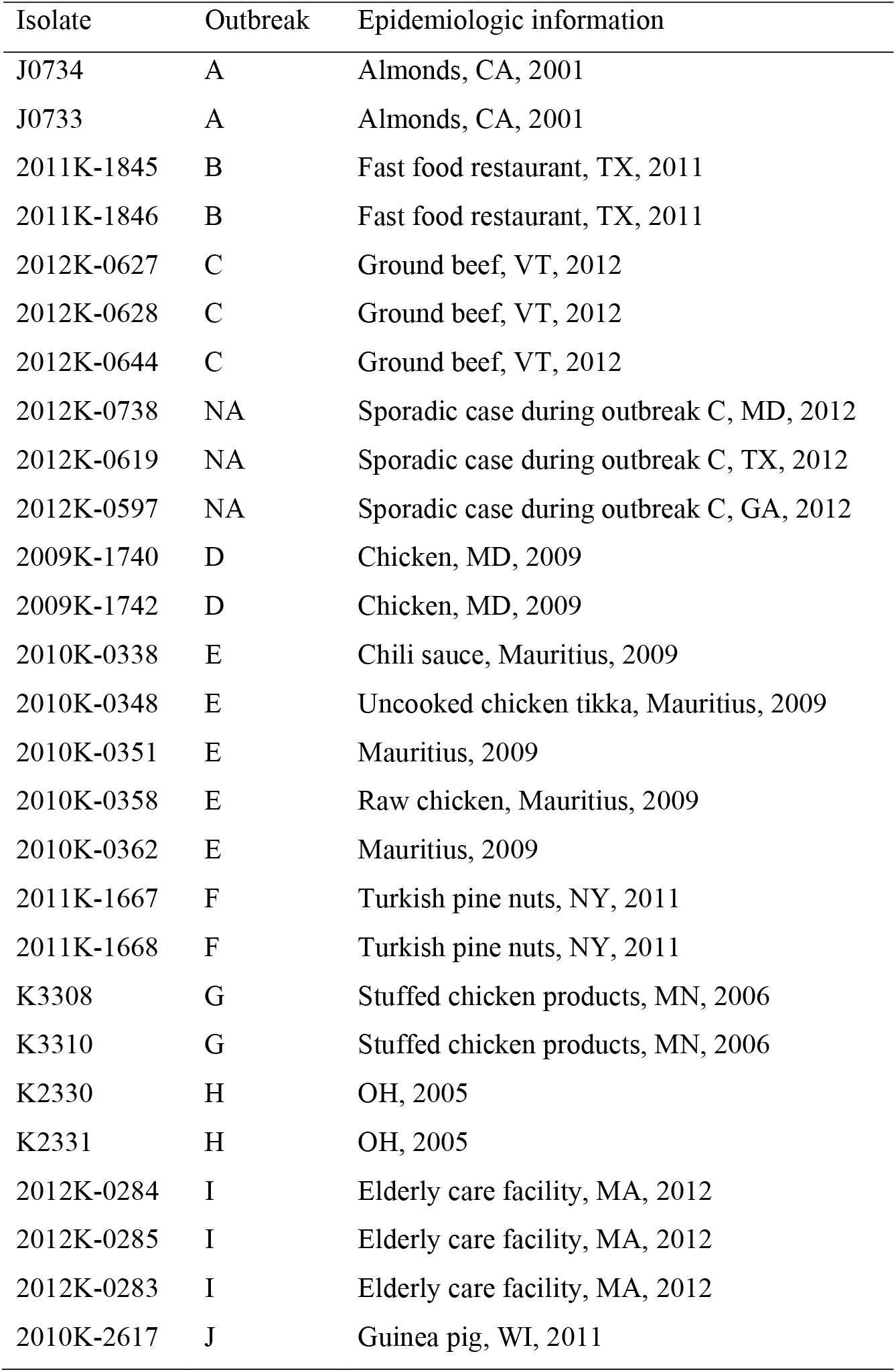
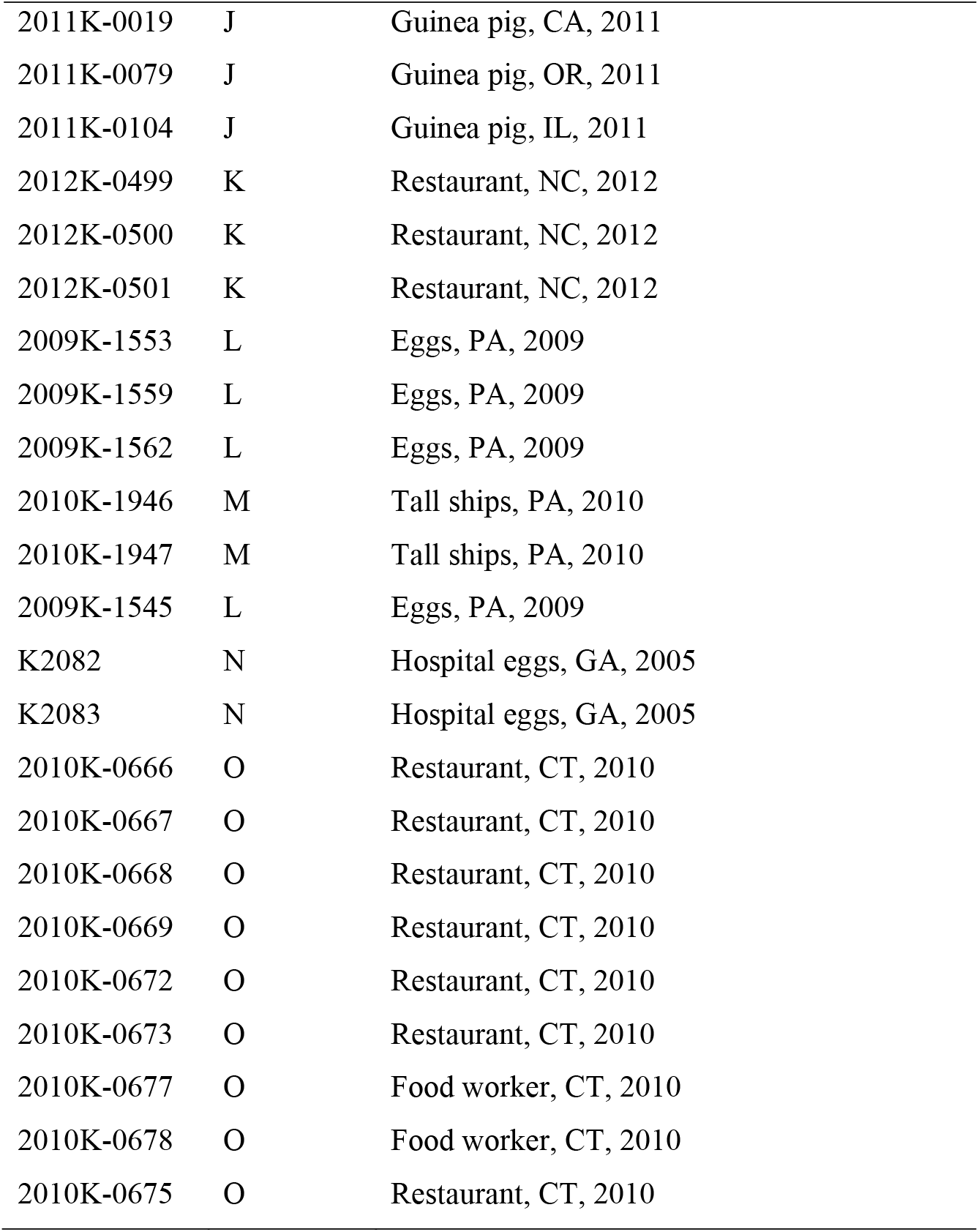
Isolates used in this study.

| Isolate | Outbreak | Epidemiologic information |
| --- | --- | --- |
| J0734 | A | Almonds, CA, 2001 |
| J0733 | A | Almonds, CA, 2001 |
| 2011K-1845 | B | Fast food restaurant, TX, 2011 |
| 2011K-1846 | B | Fast food restaurant, TX, 2011 |
| 2012K-0627 | C | Ground beef, VT, 2012 |
| 2012K-0628 | C | Ground beef, VT, 2012 |
| 2012K-0644 | C | Ground beef, VT, 2012 |
| 2012K-0738 | NA | Sporadic case during outbreak C, MD, 2012 |
| 2012K-0619 | NA | Sporadic case during outbreak C, TX, 2012 |
| 2012K-0597 | NA | Sporadic case during outbreak C, GA, 2012 |
| 2009K-1740 | D | Chicken, MD, 2009 |
| 2009K-1742 | D | Chicken, MD, 2009 |
| 2010K-0338 | E | Chili sauce, Mauritius, 2009 |
| 2010K-0348 | E | Uncooked chicken tikka, Mauritius, 2009 |
| 2010K-0351 | E | Mauritius, 2009 |
| 2010K-0358 | E | Raw chicken, Mauritius, 2009 |
| 2010K-0362 | E | Mauritius, 2009 |
| 2011K-1667 | F | Turkish pine nuts, NY, 2011 |
| 2011K-1668 | F | Turkish pine nuts, NY, 2011 |
| K3308 | G | Stuffed chicken products, MN, 2006 |
| K3310 | G | Stuffed chicken products, MN, 2006 |
| K2330 | H | OH, 2005 |
| K2331 | H | OH, 2005 |
| 2012K-0284 | I | Elderly care facility, MA, 2012 |
| 2012K-0285 | I | Elderly care facility, MA, 2012 |
| 2012K-0283 | I | Elderly care facility, MA, 2012 |
| 2010K-2617 | J | Guinea pig, WI, 2011 |
| 2011K-0019 | J | Guinea pig, CA, 2011 |
| 2011K-0079 | J | Guinea pig, OR, 2011 |
| 2011K-0104 | J | Guinea pig, IL, 2011 |
| 2012K-0499 | K | Restaurant, NC, 2012 |
| 2012K-0500 | K | Restaurant, NC, 2012 |
| 2012K-0501 | K | Restaurant, NC, 2012 |
| 2009K-1553 | L | Eggs, PA, 2009 |
| 2009K-1559 | L | Eggs, PA, 2009 |
| 2009K-1562 | L | Eggs, PA, 2009 |
| 2010K-1946 | M | Tall ships, PA, 2010 |
| 2010K-1947 | M | Tall ships, PA, 2010 |
| 2009K-1545 | L | Eggs, PA, 2009 |
| K2082 | N | Hospital eggs, GA, 2005 |
| K2083 | N | Hospital eggs, GA, 2005 |
| 2010K-0666 | O | Restaurant, CT, 2010 |
| 2010K-0667 | O | Restaurant, CT, 2010 |
| 2010K-0668 | O | Restaurant, CT, 2010 |
| 2010K-0669 | O | Restaurant, CT, 2010 |
| 2010K-0672 | O | Restaurant, CT, 2010 |
| 2010K-0673 | O | Restaurant, CT, 2010 |
| 2010K-0677 | O | Food worker, CT, 2010 |
| 2010K-0678 | O | Food worker, CT, 2010 |
| 2010K-0675 | O | Restaurant, CT, 2010 |

### WGS

Bacterial strains were grown in Tryptic Soy broth at 37 °C for 24 h. DNA extraction was performed using DNeasy Blood and Tissue Kit (Qiagen), followed by purification using AMPure XP beads (Beckman Coulter). Sample DNA concentrations were determined using a Qubit BR dsDNA assay kit (Invitrogen). For Illumina sequencing, libraries were prepared using Nextera XT DNA library prep kit and Nextera XT index kit (Illumina), and sequenced on a NextSeq 2000 instrument using NextSeq P2 reagents (Illumina) to generate 150-cycle paired end reads. For ONT sequencing, libraries were prepared using Rapid Barcoding kit (SQK-RBK004, Oxford Nanopore) and sequenced for 48 h using R9.4.1 flow cells (FLO-MIN106, Oxford Nanopore) on a GridION instrument equipped with MinKNOW software 21.05.12 (MinKNOW Core 4.37 and Bream 6.2.5, Nanopore Oxford). The isolates in sets of three were barcoded and pooled for sequencing on a single flow cell. Basecalling was performed using Guppy 5.0.12 (Oxford Nanopore) with the “super accurate” basecall model. The trim-barcodes option was selected with a minimum score of 60. The minimum qscore set for reads filtering was 10.

### Reads processing and genome assembly

For Illumina reads, Trimmomatic 0.40 (21) was used to remove adaptors and leading and trailing low quality bases. SPAdes 3.14.1 (22) was used to assemble the reads into draft genomes using the “--careful” option. For ONT reads, Porechop 0.2.4 (23) was used to remove adaptor sequences from the reads. Filtlong 0.2.0 (https://github.com/rrwick/Filtlong) was used to remove reads shorter than 1000 bp and select 1 Gb best quality data (200X depth) for downstream analyses. The selected reads were assembled using Flye 2.8.3 (24), followed by one round of polishing by Rebaler 0.2.0 (23) and two rounds of polishing by Medaka 1.4.3 (https://github.com/nanoporetech/medaka). To improve the accuracy of assembled genomes, a hybrid assembly approach that combines Illumina and ONT reads was taken using Unicycler 0.4.9 (25). Hybrid assemblies were used as the references to evaluate genome assemblies by Illumina and ONT reads alone by QUAST 5.0.2 (26). For an assembly, its error rate was calculated as the total occurrences of mismatches and indel errors per 100 Kb sequence.

### SNP typing

For Illumina reads, SNP calling was performed by CFSAN SNP pipeline 2.2.1 (27) using default settings with P125109 genome (NCBI accession AM933172.1) as the reference. For ONT reads that were selected by Filtong, SNP calling was performed through modifying the CFSAN SNP pipeline by replacing Bowtie2 with the “ont2d” option of BWA-MEM (28) for reads mapping against the P125109 reference genome. SNPs called from Illumina or ONT reads were used to build maximum-likelihood phylogenies using RAxML 8.2.12 (29). To compare SNP phylogenies between Illumina and ONT data, a tanglegram was created using R package dendextend (https://cran.r-project.org/web/packages/dendextend/vignettes/dendextend.html), which connects corresponding isolates in two juxtaposed phylogenies by minimizing intersection. The core genome of the analyzed cohort was defined as any base in the reference genome that was conserved among all the genomes in the cohort. Specifically, any base in the reference genome that was mapped by at least seven reads of an isolate genome was considered conserved in the isolate. The sensitivity and specificity of detecting SNP sites in the core genome by ONT reads was calculated by considering SNP sites identified by Illumina reads as true positives (TP), by Illumina reads but not ONT reads as false negatives (FN), and by ONT reads but not Illumina reads as false positives (FP). Non-SNP sites in the core genome identified by Illumina reads were considered true negatives (TN). Sensitivity and specificity was computed as true positive rate (TP/TP+FN) and true negative rate (TN/TN+FP).

### Homopolymer error correction (HEC)

A genome-wide HEC pipeline (gHEC) was developed to correct homopolymer errors in ONT reads throughout the 3,002 cgMLST loci of *Salmonella* defined by EnteroBase (30). gHEC was designed under the same assumption as NanoMLST (18) that if a homopolymeric tract (HT) is present at a locus, its correct size is likely the most common size of all the archived allelic HTs at the locus. Briefly, potentially low-quality alleles in the EnteroBase *Salmonella* cgMLST scheme were detected by the “PrepExternalSchema” function of chewBBACA (31) and removed from the scheme. A total of 267,108 alleles (4.4% of all alleles in the scheme), including all alleles of two loci (STMMW_09661 and STMMW_11461), were removed. These alleles contained disrupted open reading frames, absence of a stop codon, or presence of a premature stop codon. The remaining alleles were aligned against each genome assembled from ONT reads using BLAST 2.11.0 (32). For any locus where none of the archived alleles aligned perfectly with the genome, up to 15 top alignments that had a maximum of one mismatch were selected and scanned for HTs of different sizes in the aligned alleles. If the majority HT size among the allele was different from that at the locus in the genome, the HT in the genome was replaced by the most common version of the HT.

### cgMLST

EnteroBase Tool Kit (EToki) (https://github.com/zheminzhou/EToKi) was used to determine the allelic profile of each genome assembly according to the 3,002-loci cgMLST scheme for *Salmonella*. For cgMLST of genomes assembled from ONT reads, HEC was performed, followed by allelic calling as previously described. Allelic profiles were used to construct minimum spanning trees (MSTs) through GrapeTree 1.5.0 (33). Allelic differences (ADs) were calculated for all pairs of two isolates in the data set (1,225 pairs) and for pairs within each outbreak cluster. One-way analysis of variance (ANOVA) followed by Tukey HSD test was used to determine whether there were statistically significant differences between means of ADs. cgMLST by hybrid assemblies was used as the benchmark to evaluate cgMLST by ONT or Illumina reads.

### Effect of sequencing coverage on SNP and cgMLST analyses

Ten genomes each randomly selected from a distinct outbreak or a sporadic case (2010K-0673, 2011K-1667, 2011K-1845, 2012K-0499, 2012K-0644, 2012K-0738, J0734, K2083, K2331 and K3310) were used to evaluate how sequencing coverage affected SNP and cgMLST allelic calling. For each of these genomes, sequencing reads achieving the average coverage of 10X, 20X, 30X, 50X, 75X, 100X and 200X across the genome were extracted for SNP and cgMLST analyses as previously described, including applying HEC for cgMLST. The numbers of SNP sites identified and the ADs between ONT and hybrid assemblies were plotted against the sequencing coverages.

### Sequencing data and software availability

All sequencing data were deposited in the NCBI Sequence Read Archive (http://www.ncbi.nlm.nih.gov/bioproject) under project number PRJNAXXX. The gHEC pipeline is available at https://github.com/denglab/cgmlst_hec.

## RESULTS

### Base calling accuracy

The base calling accuracies of ONT and Illumina reads were evaluated by mapping the reads to their corresponding hybrid assemblies. Mismatches, insertions and deletions (indels), and overall error rates were averaged among the 50 genomes of outbreak or sporadic case isolates and summarized in Table 2. The average occurrence of indel errors per 100 Kb by ONT reads was 7.5 times as high as that by Illumina reads, with the bulk of the higher errors attributed to misidentification of HT sizes.

**Table 2.**
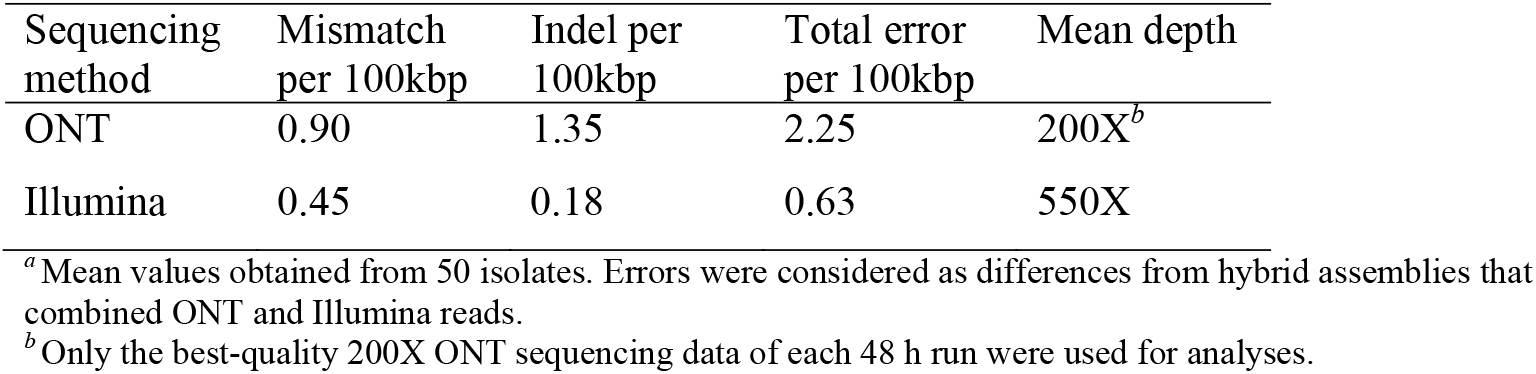
Overall sequencing accuracies^*a*^.

### SNP subtyping

A total of 2,361 and 2,365 core genome SNP sites were identified from all the isolates, including the reference genome (P125109), using Illumina and ONT reads, respectively. Among these SNP sites, 2,299 were conserved between both types of reads, 62 were unique to Illumina reads, and 66 were unique to ONT reads. Benchmarked by SNP sites identified by Illumina reads, ONT reads achieved 0.97 sensitivity and 1.0 specificity in detecting core SNP sites from the core genome of the 51 isolates (core genome size = 4,555,219 bp, 97.2% of the reference genome).

Phylogenetic trees based on core genome SNPs called from Illumina and ONT reads were largely concordant with each other (Figure 1a). Each of the 15 outbreaks was correctly resolved by both trees, with all but one isolate (2009K-1545) falling into their respective outbreak clusters, which was consistent with a previous study (20). Pairwise SNP distances between isolates were similarly concordant between Illumina and ONT reads. Linear regression analysis shows highly consistent pairwise SNP distances measured by Illumina and ONT reads (Figure 1b), indicated by a small intercept of −0.0605, a slope close to 1, and a perfect goodness-of-fit (R^2^=1.00). When placed on the same core genome SNP phylogeny (Figure 1c), ONT and Illumina sequenced genomes of the same isolate always clustered together, indistinguishably (47 isolates) or with only 1 SNP apart (3 isolates).

**Figure 1.**
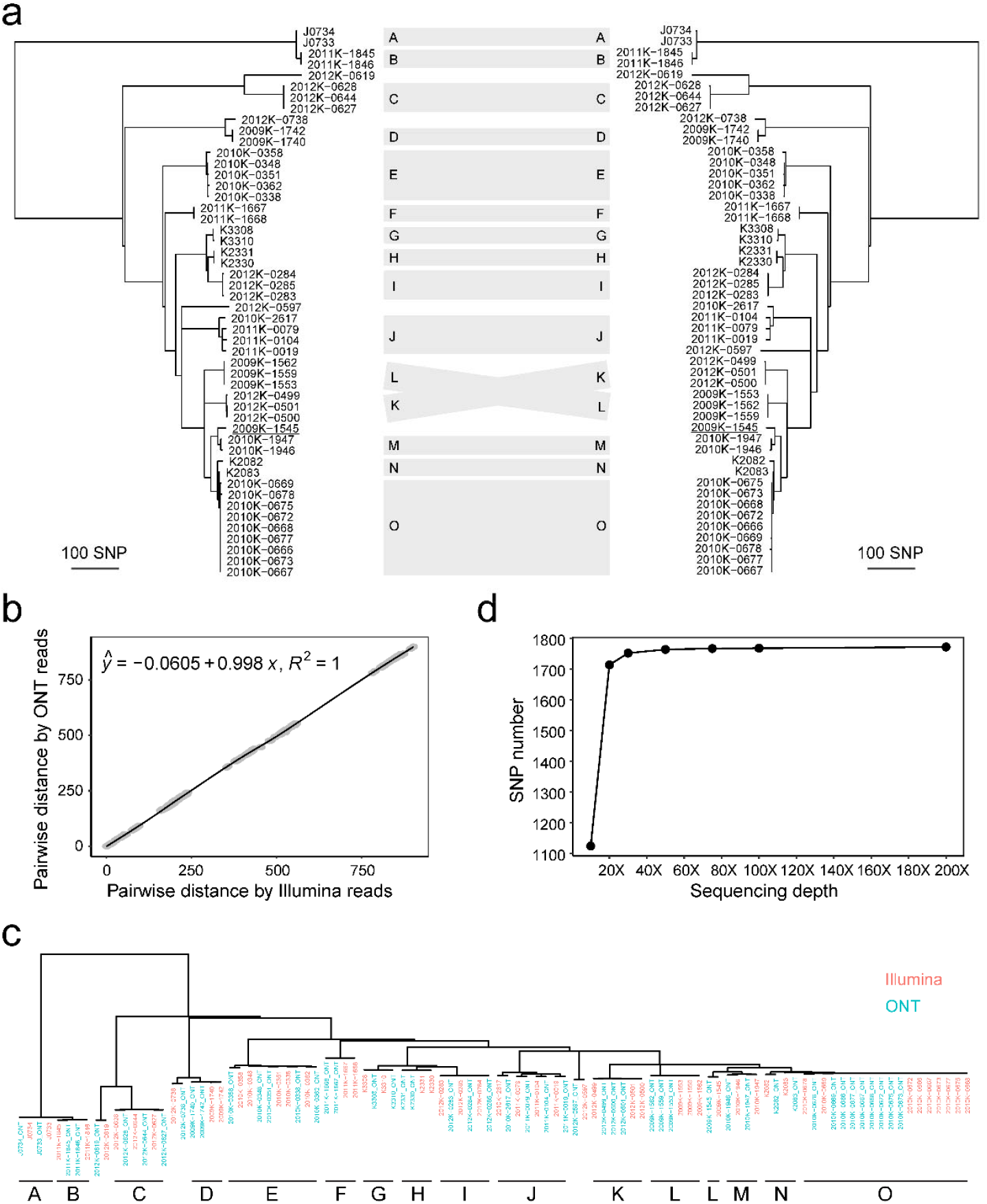
SNP subtyping. **a** SNP phylogenies constructed from Illumina (left) and ONT (right) reads. Scale bars indicate 10 SNPs. Outbreak clusters are labeled by A through O. 2009K-1545 (underlined) is the only isolate that does not cluster with other isolates from the same outbreak (Outbreak L). **b** Linear regression analysis between pairwise SNP distances (n=1,225) measured by Illumina and ONT reads. **c** SNP phylogeny constructed by co-analyzing Illumina reads (orange color) and ONT reads (cyan color). **d** Numbers of SNP sites detected by ONT reads at different sequencing depths. At the sequencing depth of 200X, reads from each 48 h run were collected from high to low quality until an average depth of 200X across the genome was reached. Reads at other depths were retrospectively sampled at different times during the run for each of the 10 selected isolates.

We estimated how ONT sequencing depths affected SNP detection (Figure 1d). ONT reads achieving the average sequencing depth of 10X, 20X, 50X, 100X and 200X for each of the analyzed genomes allowed 63.3%, 96.5%, 99.3%, 99.5% and 99.7% SNPs to be detected, respectively. By multiplexing three isolates on one flow cell, the 50X depth was achieved between 1.7 and 3.2 hours of sequencing (2.3 hours in average).

### cgMLST

Benchmarked by allelic calls from hybrid assemblies, Illumina and ONT reads averaged 1.5 and 16.4 allelic call errors (including missing allelic calls) per isolate (Figure 2, Table S1), respectively. All the allelic call errors from Illumina reads were likely caused by gaps in genome assemblies. Most of the allelic call errors from ONT reads were attributed to homopolymer errors (89.5%), with the rest being mismatches (3.2%) and insertions (7.2%) (Figure S1).

**Figure 2.**
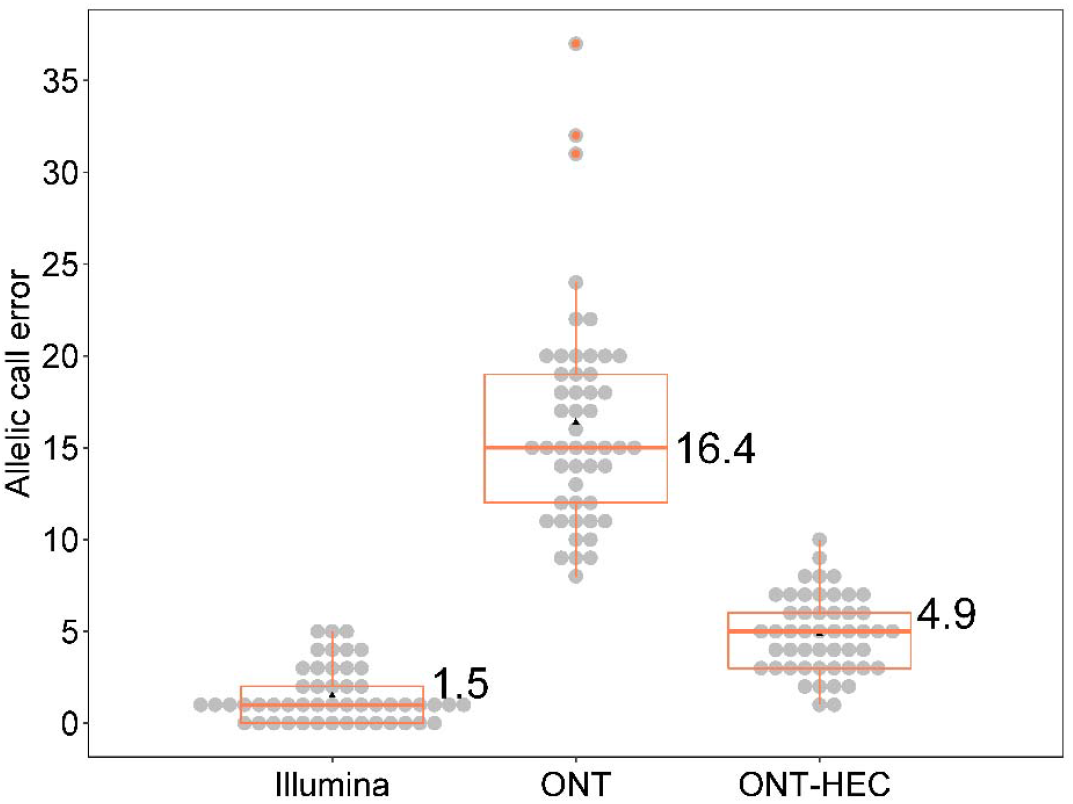
Comparison of allelic call errors among different types of reads. For each of the 50 genomes, the allelic call error is defined as the allelic difference between the cgMLST profile determined by a particular type of reads (Illumina, ONT or ONT after homopolymer error correction) and that by the hybrid assembly of the genome. The mean allelic error number for each type of reads is shown. According to one-way ANOVA followed by the Tukey HSD test, the means are significantly different from each other (P < 0.01).

MSTs based on cgMLST using hybrid assemblies (MST-hybrid, Figure 3a), Illumina reads (MST-Illumina, Figure 3b), or ONT reads (MST-ONT, Figure 3c) correctly resolved each outbreak, showing expected clustering of isolates by outbreak association. The majority of outbreak clusters exhibit longer intra-cluster branches on MST-ONT (Figure 3c) than on MST-hybrid (Figure 3a) and MST-Illumina (Figure 3b), suggesting overestimated ADs between outbreak isolates by ONT reads largely due to homopolymer size misidentification.

**Figure 3.**
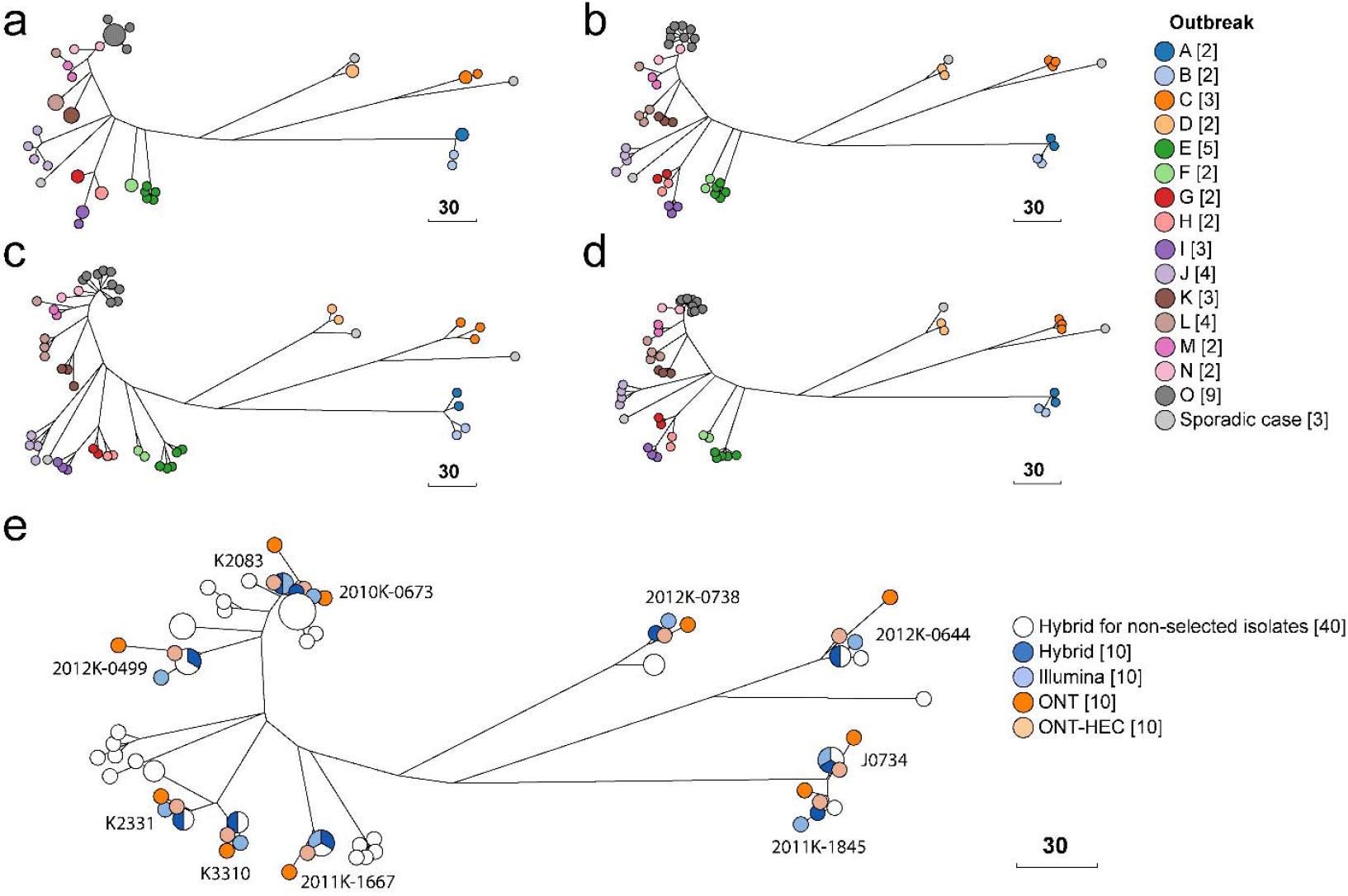
Minimum spanning trees (MSTs) constructed by cgMLST. **a** MST by hybrid assemblies. **b** MST by Illumina reads. **c** MST by ONT reads. **d** MST by ONT reads after homopolymer error correction. **e** MST by cgMLST combining hybrid assemblies, Illumina reads, ONT reads, and ONT reads after homopolymer error correction for 10 representative isolates. Scale bars indicate 30 allele differences.

We adopted an empirical strategy for HEC originally designed for seven-gene MLST (18) and expanded it to the 3,002-gene scheme. The HEC workflow started with identification and removal of low-quality alleles (see Materials and Methods). Genome-wide HEC corrected 94.8% of the homopolymer errors, reducing average allelic call errors from 16.4 to 4.8 errors per isolate (Figure 2, Table S1). In comparison, only 11.0% allelic call errors were corrected using the EnteroBase cgMLST database without the filtering of low quality alleles. This was because the vast majority of ONT-sequenced HTs, regardless of their sizes having been correctly resolved or not, matched exactly with archived alleles in EnteroBase. This observation suggested that alleles with the same homopolymer errors had already been deposited into EnteroBase. Further outcome assessment of the genome-wide HEC (Table S2) showed that 695 of the 733 homopolymer errors that occurred across 79 loci among the 50 ONT sequenced isolates had been successfully corrected, including complete clearance of 68 loci of any homopolymer errors. Six loci (STMMW_05601, STMMW_30521, STMMW_15211, STMMW_19051, STMMW_19211 and STMMW_27051) with 24 homopolymer errors escaped HEC because the errors happened to cause exact matches to other alleles. HEC also failed to correct 14 homopolymer errors on another five loci (STMMW_28541, STMMW_11241, STMMW_38201, STMMW_13711 and STMMW_06641). Interestingly, HEC introduced 116 new allelic call errors to 14 loci, including two loci (STMMW_39811 and STM4351) where 46 of the 50 isolates were affected (Table S3). Inspection of these loci showed that the miscorrected alleles, on which uncorrected ONT reads had originally agreed with Illumina and hybrid assemblies, had issues such as incomplete or disrupted reading frames. The HEC workflow precluded such alleles, forcing the selection of other alleles as the basis for correction, which led to the miscorrections. This finding suggests the presence of alleles with potential or perceived sequence abnormalities outside HTs in EnteroBase. Despite HEC escapees and miscorrections, HEC achieved a net reduction of 577 homopolymer errors, reducing homopolymer error prevalence from 14.7 to 0.8 occurrences per isolate. After HEC, the remaining allelic call errors from ONT reads (n=242) were attributed to homopolymer errors (15.7), the rest being mismatches (10.7%), insertions (24.4) and miscorrections (48.8%) (Figure S1). cgMLST using post-HEC ONT reads led to tighter clustering of isolates within multiple outbreak clusters on MST-ONT (Figure 3d). Mixed use of post-HEC ONT reads and Illumina reads in cgMLST achieved concordant results between the two types of reads, as shown by the tight clustering of Illumina- and ONT-sequenced samples of the same isolate (Figure 3e).

The application of HEC substantially improved the estimation of cgMLST pairwise distances between ONT sequenced isolates (Figure 4) for ONT sequenced isolates. Prior to HEC, most of the pairwise distances estimated by ONT reads exceeded those by hybrid assemblies by more than 10 ADs (Figure 4a). HEC removed this systematic inflation and curbed the vast majority of remaining discrepancies within five ADs (Figure 4b). At certain loci in some isolates, originally different allelic calls between isolates were incorrectly changed to the same calls by HEC. This convergence caused by HEC errors led to certain underestimated ADs by ONT reads (blue cells, Figure 4b).

**Figure 4.**
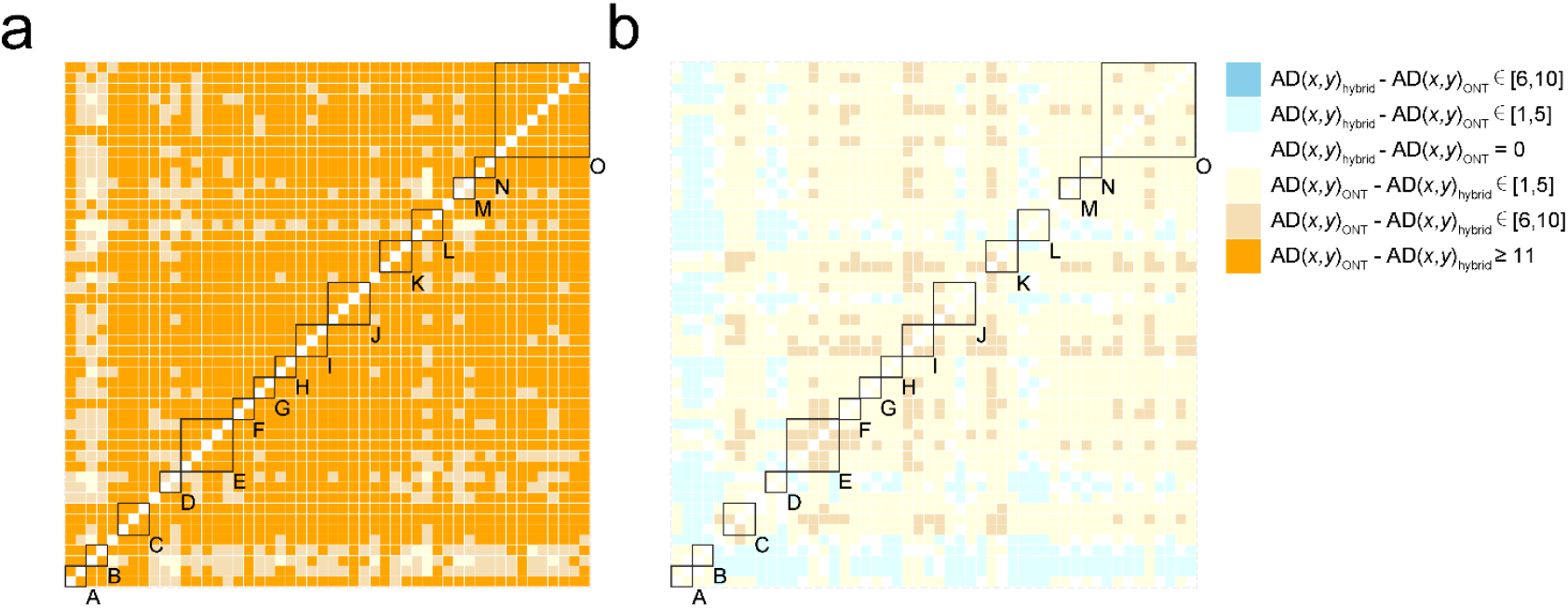
Effect of Homopolymer error correction (HEC) on cgMLST pairwise distance between isolates estimated by ONT reads. Define AD(*x,y*)_ONT_ as the allelic difference between isolate *x* and isolate *y* using ONT reads, and AD(*x,y*)_hybrid_ as that by hybrid assemblies. **a** A heatmap summarizing the difference between AD(*x,y*)_ONT_ and AD(*x,y*)_hybrid_ for each of the 1,225 combinations of *x* and *y* before HEC. **b** A heatmap summarizing the difference between AD(*x,y*)_ONT_ and AD(*x,y*)_hybrid_ for each of the 1,225 combinations of *x* and *y* after HEC.

For isolates within individual outbreak clusters, the maximum pairwise distances measured by ONT reads without HEC were significantly different from those by hybrid assemblies and Illumina reads (Table 3). After HEC, cgMLST by ONT reads performed statistically similar to those by hybrid assemblies and Illumina reads in estimating intra-outbreak ADs (Table 3). According to hybrid assemblies, isolates in 13 of the 15 outbreak clusters belonged to the same HC10 group (pairwise difference < 10 ADs) using hierarchical clustering of cgMLST (HierCC) (30). Without HEC, the HC10 grouping was violated in 11 of the 13 outbreak clusters due to excessive homopolymer errors (pairwise difference > 10 ADs, Table 3). After HEC, the HC10 grouping was restored in 10 of the 11 cluster (Table 3, shaded rows). For Outbreak E isolates, homopolymer errors caused a maximum AD of 24, compared to 4 by hybrid assemblies. HEC reduced the maximum AD to 11 but still failed to restore its HC10 grouping (Table 3).

**Table 3.** Maximum ADs for isolates within each outbreak.

| Outbreak<sup>#</sup> | Number of isolates | Hybrid assemblies | Illumina reads | ONT reads | ONT reads after HEC |
| --- | --- | --- | --- | --- | --- |
| A | 2 | 0 | 1 | 16 | 2 |
| B | 2 | 1 | 6 | 14 | 2 |
| C | 3 | 1 | 6 | 34 | 8 |
| D | 2 | 0 | 3 | 8 | 2 |
| E | 5 | 4 | 4 | 24 | 11 |
| F | 2 | 0 | 1 | 22 | 5 |
| G | 2 | 0 | 2 | 15 | 3 |
| H | 2 | 0 | 2 | 15 | 1 |
| I | 3 | 1 | 10 | 23 | 7 |
| J | 4 | 20 | 25 | 39 | 24 |
| K | 3 | 0 | 5 | 26 | 6 |
| L* | 3 | 0 | 5 | 15 | 2 |
| M | 2 | 2 | 3 | 9 | 3 |
| N | 2 | 11 | 12 | 27 | 14 |
| O | 9 | 3 | 10 | 23 | 7 |
| Mean | 3.1 | 2.9<sup>a</sup> | 6.3<sup>a</sup> | 20.7<sup>b</sup> | 6.5<sup>a</sup> |

<sup>#</sup> Shaded rows indicate outbreak clusters whose original HC10 grouping (by hybrid assembly) was violated by ONT reads but restored after HEC.
\* 2009K-1545 was excluded from Outbreak L as it was phylogenetically distinct from other outbreak isolates
<sup>a</sup> One-way ANOVA followed by the Tukey HSD test showed no significant difference between the means of these groups ( $p > 0.01$ ).
<sup>b</sup> One-way ANOVA followed by the Tukey HSD test showed significant difference between the means of this group and those of the others ( $p < 0.01$ ).

We estimated how sequencing depths by ONT affected cgMLST through counting allelic call errors against hybrid assemblies at different depths. At 100X depth, the mean allelic call errors of the 10 representative isolates was statistically similar to that at 200X depth (Figure 5). By multiplexing three isolates on one flow cell, 100X depth was attained between 3.2 and 6.6 hours of sequencing (4.6 hours in average).

**Figure 5.**
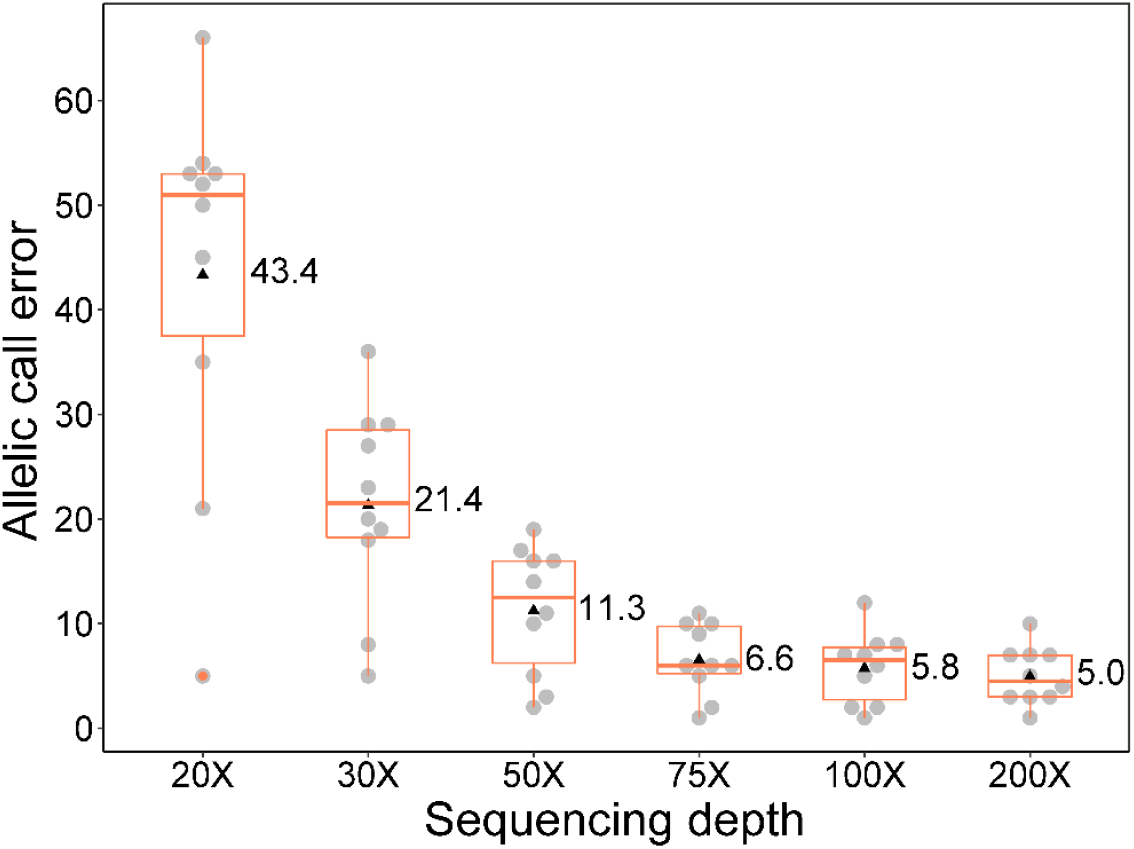
cgMLST allelic call errors at different sequencing depths using ONT. For each of the 10 representative genomes, the allelic call error is defined as the allelic difference (AD) between ONT reads- and hybrid assembly-determined cgMLST sequence profiles of the genome. ONT reads were pre-processed by homopolymer error correction. The mean allelic error number for each depth is shown. At the sequencing depth of 200X, reads from each 48 h run were collected from high to low quality until an average depth of 200X across the genome was reached. Reads at other depths were retrospectively sampled at different times during the run. According to one-way ANOVA followed by the Tukey HSD test, the mean allelic error numbers at 50X, 75X, 100X and 200X for each type are not significantly different from each other (P < 0.01).

## DISCUSSION

Continuous iteration and evolution of the nanopore technology, including both sequencing chemistry and analytical practices, has posed challenges to its evaluation and standardization for routine applications in public health and food industry. In this study, we evaluated R9.4.1 flow cells, which have matured into a mainstream option of WGS by ONT. Using the state-of-art bioinformatics workflows, our evaluation suggests that the ONT platform has reached technical readiness acceptable for strain-level subtyping of common foodborne pathogens such as *S. enterica*.

SNP detection that targets a single base difference flanked by conserved sequences is inherently insusceptible to homopolymer size misidentifications that constitute a major source of base calling errors in ONT reads. As expected, ONT reads delivered SNP calls highly consistent with those by Illumina reads, allowing accurate phylogenetic placement and distance estimation among closely related *Salmonella* Enteritidis isolates such as those from monoclonal outbreaks. Should WGS by ONT be implemented for foodborne pathogen surveillance, a likely scenario is the mixed use of ONT and Illumina reads in subtyping analyses. By combining both types of reads, we observed zero (n=47) or one (n=3) SNP discrepancy between ONT- and Illumina-sequenced genomes of the same isolate. This was improved from six or seven SNP discrepancies between Illumina and ONT sequences similarly measured with *E. coli* isolates (15), which may be explained by newer bioinformatics tools and/or flow cells used in the present study. Similarly, combining Illumina and post-HEC ONT reads yielded concordant clustering of isolates using cgMLST between the two types of reads. These results suggest that ONT and Illumina reads can be used individually or together for strain-level subtyping of bacterial isolates.

In contrast to SNP subtyping, cgMLST requires accurate allelic calls for thousands of predefined loci across the genome. This is considerably challenging given the homopolymer error rates affecting the base calling accuracy of ONT reads. With R9.4.1 reads, we found that the HEC approach effectively ameliorated homopolymer errors, reducing the average incorrect allelic calls from 16.4 to 4.8 per isolate (14.7 to 0.8 per isolate for homopolymer errors). With fewer than five errors across 3,002 loci, the average allelic calling accuracy was 99.8%. Epidemic outbreaks of *Salmonella* are often associated with hierarchical clusters of cgMLST (HierCC) defined by empirical cutoffs of 2, 5 or 10 intra-cluster ADs (HC2, HC5 or HC10) (30). HEC is therefore useful to restore the close relatedness between isolates to trigger outbreak investigation (e.g. HC10), which could be masked by homopolymer errors in ONT reads. In our evaluation, HEC was able to restore the HC10 grouping for 10 out of 11 outbreak clusters (Table 3), allowing ONT reads to perform statistically similarly to Illumina reads in grouping closely related outbreak isolates. Notably, HEC can introduce miscorrections. The HEC procedure is initiated by a lack of exact match between a query sequence and any of the quality-filtered alleles in EnteroBase. Using the allele quality criteria by Chewbacca (31), the workflow precludes any allele with an incomplete or disrupted open reading frame prior to determining the need for HEC. However, EnteroBase preserves some alleles with frameshift mutations, such as pseudogenes (34, 35) (Zhemin Zhou, personal communication). Precluding these alleles can lead to unnecessary HEC on genuine alleles without sequencing errors. Furthermore, homopolymeric tracts in prokaryotes frequently exhibit frameshift mutations and different sizes (36), thus the differentiation between genuine sequences and sequencing errors is inherently difficult. Despite these confounding factors, HEC achieved substantial net reduction of homopolymer errors to identify closely related isolates according to empirical AD thresholds for outbreak detection. Recently introduced ONT chemistry (R10.4) than the evaluated version has been reported to generate near-perfect bacterial genome assemblies using ONT reads alone (37). In particular, R10.4 showed a substantial improvement over the evaluated version (R9) in resolving homopolymer sizes, leading to correct size determination for the majority of homopolymers up to 10 bps (37). While this ability may allow HEC-free cgMLST using ONT reads, HEC can provide forward compatibility for isolates that have been or continue to be sequenced by the R9 chemistry to facilitate their co-analysis with future ONT-sequenced genomes presumably not requiring HEC. Although R10.4 lags behind R9.4.1 in theoretical output (30 Gb vs 50 Gb) due to slower translocation speed (250 bp/s by vs 400 bp/s), it is expected that further iteration(s) of R10 could match the rapid library preparation and stable output by R9.4.1 flow cells. Given the already close performance between R9.4.1 and Illumina reads in both SNP and cgMLST analyses, it remains to be determined whether R10.4 would bring incremental or significant gains for routine WGS of foodborne pathogens.

One appeal of using ONT for foodborne pathogen WGS in public health, food industry, and/or clinical settings is the rapid turnaround time, especially given the possibility of real-time analytics by ONT. A previous study suggests that multiplexing 3-5 *Salmonella* isolates on an R9.4.1 flow cell helps strike a balance between cost efficiency and sequencing output as well as minimize the impact of uneven distribution of reads among pooled isolates (13). With three isolates multiplexed on one flow cell, sufficient sequencing depth was achieved for each isolate within seven hours of sequencing to support accurate SNP typing and cgMLST analysis, suggesting a prospect of fast WGS turnaround for small batches of isolates.

In conclusion, our study established a baseline for the continuously evolving nanopore technology as a technologically viable solution to high quality subtyping of *Salmonella* using both SNP and cgMLST analyses, individually or together with Illumina platforms. The study paves way for evaluating and optimizing the logistics of implementing the ONT approach for foodborne pathogen surveillance in specific settings.

## ACKNOWLEGEMENT

The authors would like to thank Zhemin Zhou for discussion on EnteroBase and cgMLST. We are grateful for Patti Fields and Susan Van Duyne for providing isolates. We also thank Bala Ganesan, Jerome Combrisson, Aurelien Maillet, Ash Scribbins, Kristel Hauben, Becca Barnett, and John Luedke for providing review and comments.

